# Multiparametric quantitative MRI uncovers putamen microstructural changes in Parkinson’s Disease

**DOI:** 10.1101/2024.05.26.595926

**Authors:** Elior Drori, Lee Cohen, David Arkadir, Aviv A. Mezer, Gilad Yahalom

**Author notes:** These authors contributed equally to this work (Aviv A. Mezer, Gilad Yahalom).

## Abstract

Parkinson’s disease (PD) is a progressive neurodegenerative disorder dominated by motor and non-motor dysfunction. Despite extensive research, the in vivo characterization of PD-related microstructural brain changes remains an ongoing challenge, limiting advancements in diagnostic and therapeutic strategies. The putamen, a critical structure within the basal ganglia, plays a key role in regulating movement and is profoundly affected in early PD-related neurodegeneration. In this study, we collected multiparametric quantitative MRI (qMRI) brain data of PD patients and healthy controls, to investigate microstructural alterations in the putamen in PD. We utilized a gradient analysis technique to analyze the spatial variations of various qMRI parameters, including relaxation rates (R1, R2, R2*), water fraction (WF), susceptibility, magnetization transfer saturation (MTsat), and diffusion metrics (MD, FA). Our findings reveal significant spatial gradients and interhemispheric asymmetries in these biophysical properties along the anterior-posterior axis of the putamen. Notably, PD patients exhibited increased water fraction and altered transverse relaxation rate R2*, particularly in the posterior putamen, correlated with motor symptom laterality. These microstructural changes suggest underlying tissue atrophy or neuroinflammatory processes associated with PD. The study underscores the importance of the posterior putamen as a focal point for PD pathology and highlights the potential of localized gradient analysis in detecting subtle yet clinically significant brain changes. The new qMRI dataset provides valuable insights into PD pathology, potentially aiding in the development of more precise diagnostic tools and targeted therapies.

## Introduction

Parkinson’s Disease (PD) is a progressive neurodegenerative disorder primarily characterized by motor and non-motor symptoms, fundamentally altering the life quality of those affected. Despite extensive research, the microstructural neural changes associated with PD have not yet been fully characterized in vivo. This limits the ability to monitor the pathophysiological mechanisms of the disease, and consequently hinders the research and development of more effective diagnostic and therapeutic strategies^1^.

The putamen, primarily associated with the motor domain of the striatum, is a key brain structure within the basal ganglia, which plays a crucial role in regulating movement and facilitating goal-directed behavior^2^. In PD, the progressive loss of dopaminergic neurons in the substantia nigra pars compacta (SNpc) leads to a significant decrease in putaminal dopamine levels. This depletion leads to degeneration of the putamen and impairs its functionality, consequently giving rise to the characteristic motor symptoms of PD^2^.

Previous research has highlighted two spatial aspects of putamen neurodegeneration in PD. One key aspect found using PET and SPECT is a gradual dopamine depletion starting in the posterior parts and advancing anteriorly with PD progression^3,4^. Furthermore, an MRI study found that the major anatomical axis of the putamen, i.e., its anterior-posterior (AP) axis, is associated with spatial microstructural patterns that change posteriorly in early stages of PD^5^. Another key property is inter-hemispheric asymmetry; PET, SPECT and MRI studies have demonstrated asymmetry in dopamine depletion and tissue microstructure integrity, corresponding contralaterally with the asymmetrical motor signs commonly seen in early PD stages^2,4,5^.

These findings in PD are in line with the view of inherent spatial inhomogeneities of the typical striatum and particularly the putamen. Animal and postmortem human studies found molecular and connectivity gradients in the healthy striatum, using histochemical staining and neural tracing methods^6–8^. Furthermore, in vivo human studies have identified gradients of connectivity, function and biophysical properties along the major axes of the striatum, using diffusion MRI tractography, task-based functional MRI (fMRI), resting-state fMRI, and quantitative MRI (qMRI)^5,9–15^.

Recent advances in neuroimaging, particularly multiparametric qMRI, have opened new avenues for exploring the brain’s microstructure in vivo^16^. qMRI encompasses a range of imaging techniques that quantify the physical properties of brain tissue, and has been instrumental in detecting subtle biological changes in development, normal aging and neurodegenerative diseases, particularly in PD^17,18^. Key qMRI parameters include R1 (longitudinal relaxation rate), R2 (transverse relaxation rate), and R2* (effective transverse relaxation rate), which are sensitive to tissue macromolecular composition and iron content. Quantitative Susceptibility Mapping (QSM) offers information on tissue magnetism and iron deposition, while Magnetization Transfer Saturation (MTsat) indicates the exchange between free and bound protons, which depends on

the macromolecular environment. Water Fraction, another qMRI metric, helps in understanding tissue hydration and tissue density. Furthermore, Mean Diffusivity and Fractional Anisotropy, derived from Diffusion Tensor Imaging (DTI), provide information on the diffusion of water molecules, indicating the tissue structure and integrity.

In recent years, several potential disease-modifying drugs have shown promising evidence of neuroprotection^19–21^. Therefore, accurate diagnosis of Parkinson’s Disease (PD) and monitoring disease progression during studies involving these drugs are crucial for drawing meaningful conclusions about disease modification. qMRI may offer valuable insights into addressing this challenge.

In this study, we collected multiparametric qMRI data from PD patients and healthy controls (HC). This type of data is rarely available in a clinical setting due to the time-consumption and complexities involved in acquiring and analyzing it. We utilized this valuable data in conjunction with a spatial gradient analysis method developed recently^5^, providing a unique vantage point for a detailed examination of the biophysical properties of the putamen across both healthy individuals and PD patients.

We aim to unravel spatial variations in relaxation rates, WF, susceptibility, MTsat, and diffusion parameters along the AP axis of the putamen, measured on the same group of subjects. These insights are crucial for a deeper understanding of the microstructural changes in PD and their association with clinical manifestations, such as motor symptom laterality. Beside contributing to the broader understanding of PD pathology, our study also highlights the potential of localized gradient analysis in revealing subtle, yet clinically significant, deep-brain microstructural changes in neurodegenerative diseases.

## Methods

### Subjects

A total of 108 PD and HC individuals participated in the study. PD patients were diagnosed based on the Movement Disorder Society Clinical Diagnostic Criteria for Parkinson’s disease (MDS-PD Criteria). Four healthy subjects were excluded due to age mismatch, and two subjects were removed because of observed structural brain abnormalities that prevented reasonable segmentations of the regions for analysis. Not all participants completed all the imaging sessions. Visual inspection for quality assurance was conducted for each scan, and 11 subjects were excluded due to significant head motion that particularly affected the putamen region. Following this data screening process, the final analysis included 91 subjects: 62 PD patients and 29 HC subjects. The groups did not statistically significantly differ in age (*t* = 1.17, *p* = 0.24) or sex (Pearson’s *χ*^2^ = 1.73, *p* = 0.19). Demographics and PD characteristics of analyzed subjects are shown in **Table 1**. All study procedures were approved by the Helsinki Ethics Committee of Shaare Zedek Medical Center, Jerusalem, Israel. A written informed consent was obtained from all participants before the procedure.

**Table 1.** Demographics and PD characteristics. H&Y, Hoehn and Yahr scale of PD progression; MDS-UPDRS III, MDS Unified Parkinson’s Disease Rating Scale part III, assessment of motor dysfunction; MoCA, Montreal Cognitive Assessment. For further details, see Methods: Behavioral Assessments.

| Variable | PD (N=62) | HC (N=29) |
| --- | --- | --- |
| Age, years | 68.4 ± 10.1 (range, 37-86) | 70.9 ± 7.1 (range, 56-78) |
| Sex, female, N (%) | 21 (34%) | 14 (48%) |
| H&Y scale | 2.1 ± 0.8 |  |
| Disease duration, years | 5.3 ± 4.3 (range, 0-21) |  |
| MDS-UPDRS III, score | 29.8 ± 11.7 (range, 10-69) |  |
| MoCA, score | 23.2 ± 4.0 (range, 11-29) |  |

### Data Acquisition

Data were collected on a 3T Siemens MAGNETOM Skyra scanner at the Edmond and Lily Safra Center (ELSC) neuroimaging unit at the Hebrew University of Jerusalem.

#### R1, WF, R2* mapping

For quantitative R1, R2*, and WF mappings, 3D Fast Low Angle Shot (FLASH) images were acquired at four different flip angles (α = 4°, 10°, 20°, and 30°). Each image included five equally spaced echoes, with echo time (TE) of 3.34, 6.01, 8.68, 11.35, and 14.02 ms. The repetition time (TR) was 19 ms, and the scan resolution was 1 mm^3^ isotropic.

For excitation bias estimation, spin-echo inversion recovery (SEIR) images were acquired with echo-planar imaging readout (EPI). Imaging parameters included: TE = 49 ms, TR = 2920 ms, and inversion times (TI) of 200, 400, 1200, and 2400 ms. Slice thickness was 3 mm, and in-plane resolution was 2 mm^2^.

Total N=89 subjects completed the FLASH scans successfully (60 PD and 29 HC). From those, N=84 subjects also completed the SEIR scans (55 PD and 29 HC).

Whole-brain R1 and WF maps, as well as field bias maps of excitation (B1+) and receive (B1−) for correction, and a synthetic T1w contrast were computed from the FLASH and SEIR images, using mrQ software (https://github.com/mezera/mrQ/)^22,23^.

R2* maps were fitted from the FLASH scans using the MPM toolbox^24^. R2* estimates acquired from each of the four FLASH images were averaged for increased signal-to-noise ratio (SNR).

#### MTsat

For quantitative MTsat mapping, a 3D FLASH image was acquired with an additional MT pulse, at flip angle α = 10°. The image was acquired at five equally spaced echoes (TE = 3.34 to 14.02 ms) and TR = 37 ms. Scan resolution was 1 mm^3^ isotropic.

Total N=81 subjects completed the MT scans successfully (52 PD and 29 HC).

Whole-brain MTsat maps were computed from the shortest echo (3.34 ms), as described in Helms et al.^25^.

#### DTI

For DTI measurements, diffusion-weighted spin-echo echo-planar (EPI) images were acquired, with TE = 95.8 ms and TR = 6000 ms. The diffusion weighting gradients were set in 64 directions with a diffusion weighting strength of b = 2000 s/mm^2^ (G = 45mT/m, δ = 32.25 ms, Δ = 52.02 ms). Eight non-diffusion-weighted images (b = 0) were interspersed between the diffusion-weighted images. In addition, non-diffusion-weighted images with reversed phase-encoding (PE) blips were acquired. Scan resolution was 1.5 mm^3^ isotropic.

Total N=70 subjects completed the DTI scans successfully (45 PD and 25 HC).

Diffusion analysis was conducted using the FMRIB’s Diffusion Toolbox (FDT)^26^ within FSL^27^. Corrections for distortions induced by susceptibility and eddy currents were performed using the reverse PE data, utilizing the EDDY^28^ and TOPUP^29^ functions. After correction, MD and FA maps were generated using the DTIFIT function in FDT.

#### QSM

For quantitative susceptibility mapping (QSM), multi-echo gradient echo images were acquired at seven echoes (TE = 8.41, 14.04, 18.86, 22.45, 26.28, 30.89, and 35.52 ms), TR = 41 ms, and flip angle of α = 15°. Scan resolution was 1 mm^3^ isotropic.

QSM processing utilized Tikonov regularization-based estimation^30^, and using the Morphology Enabled Dipole Inversion (MEDI) toolbox^31^. This included frequency offset calculation for each voxel via complex fitting, spatial phase unwrapping, and background field removal.

Out of 63 participants who underwent susceptibility imaging, 14 were excluded from the final analysis because of substantial acquisition artifacts, resulting in a total of 49 subjects (23 PD and 26 HC).

#### R2

For quantitative T2 mapping, multi‐echo spin echo images were acquired at 10 equally spaced echoes between 12 ms and 120 ms, and TR = 4210 ms. Scan resolution was 2 mm^3^ isotropic.

Whole-brain R2 (1/T2) maps were fitted using the nine longest TEs (shortest TE = 12 ms was excluded), by solving the following T2-weighted signal (S) equation^32^, simultaneously for T2 and M0:

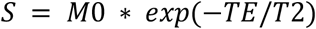

Total N=62 subjects completed the T2 scans successfully (34 PD and 28 HC).

#### Behavioral Assessments

##### Motor function and asymmetry

PD patients’ motor signs were assessed through the Movement Disorder Society-Unified Parkinson’s Disease Rating Scale (MDS-UPDRS-III)^33^. The patients’ scores were within the range of 10 to 69, with a mean ± SD of 29.8 ± 11.7. Hoen and Yahr stages were 2.1 ± 0.8. Three patients did not have the UPDRS data and were excluded from related analyses. The motor signs asymmetry score of each individual was calculated by subtracting the total score of right body-side items from the left body-side items (L-R). A higher positive score indicates left-side predominance, while a higher negative score indicates right-side predominance.

For patients exhibiting left-side predominance, the Most Affected Side (MAS) was identified as the left side of the body and the contralateral (right) putamen, while the Less Affected Side (LAS) was identified as the right side of the body and the left putamen. Conversely, for those with right-side predominance, the MAS and LAS definitions were reversed. Patients displaying no motor asymmetry were not included in analyses that compared MAS with LAS.

##### Cognitive assessments

The cognitive condition of PD patients was evaluated using the Montreal Cognitive Assessment (MoCA)^34^. Two patients did not have MoCA data. Scores ranged from 11 to 29, with mean ± SD of 23.2 ± 4.0. Among the patients, 35% fell within normal range (scores between 26-30), 55% exhibited mild cognitive impairment (scores between 18-25), and 10% were identified as having moderate cognitive impairment (scores between 10-17). No patients were found to have severe cognitive impairment (scores under 10).

### Data Analysis

#### Regions of Interest (ROIs)

Putamen ROIs were delineated using FSL’s FIRST probabilistic segmentation tool^35^. The subject’s synthetic T1w image was used as a reference image. To minimize partial volume effects, the outer 1-mm layer of the ROIs were removed. Visual inspection of all segmented ROIs was conducted to ensure quality. The segmentation was transformed also to the R2 space (2 mm^3^) and the DTI space (1.5 mm^3^). For R2, a rigid body transformation from R1 to R2 was computed using SPM (www.fil.ion.ucl.ac.uk/spm/software/spm12/) and was then applied on the segmentation volume. In the case of DTI, a nonlinear transformation from R1 to FA was performed using ANTS^36^, and was then applied on the segmentation.

#### Putamen AP Gradients

We computed qMRI gradients along the putamen’s major (AP) axis as described in Drori et al.^5^, using the mrGrad toolbox (https://github.com/MezerLab/mrGrad/). In short, the major axis of the putamen structure is computed using singular value decomposition (SVD) of the ROI coordinates, independently of the image axis system and image intensities. The image values (i.e., the qMRI parameter) is then sampled along the axis, as the median value of each equally-spaced segment along the axis. In our study we used 20 equally spaced segments along the AP axis. The obtained values represent a spatial function of the sampled parameter along the axis. We then smoothed the profile using a moving gaussian kernel of size 5.

#### Interhemispheric asymmetry

We defined interhemispheric asymmetry on the subject level as the left-hemisphere value (in an ROI or a gradient node) minus the right-hemisphere value. With this definition, a negative value reflects a higher value in the right hemisphere, and a positive value reflects a higher value in the left hemisphere.

#### Statistical analysis

##### Linear mixed-effects statistical models

Statistical inference for spatial and disease-related effects was performed using linear mixed-effects models (LMMs). For each MRI metric, the position along the putamen AP axis (twenty ordinal levels from 0 to 1), and the clinical group (two categorical levels: PD, HC), were included as fixed effects. Subject ID was included as a random effect, and the MRI metric served as the dependent variable. At first, we also included Age and Sex as covariates for the Clinical Group:

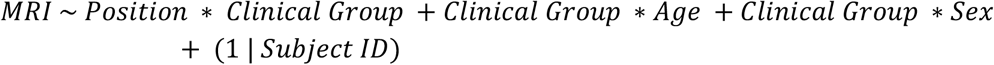

As no interactions were found between Clinical Group and either Age or Sex, in further analyses we excluded these variables from the model:

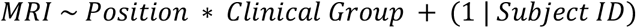

For evaluation of quadratic spatial effects, we also integrated a quadratic term for the position:

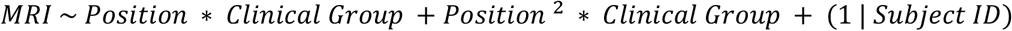

Standardized beta coefficients (*β)* were computed by z-scoring all continuous variables, including Age and the MRI response variables. Position was not z-scored as it was already normalized between 0 and 1, allowing *β*_*Position*_ to represent the estimated change along the entire AP axis of the putamen.

#### Post-hoc tests between groups

In cases where significant interactions between Position and Clinical Group were detected in the models, post hoc analyses were conducted using two-sample t-tests between PD and HC groups at each positional level along the axis. Given the dependence among these comparisons, we applied corrections for multiple comparisons using the Nichols and Holmes permutation test^37^.

#### PD-related variable correlations

To test the relationships between the putamen’s gradient asymmetry and the motor function asymmetry, we fitted linear regression models. We report R^2^ values, representing the proportion of variance explained by the structural asymmetry variable. To rule out effects of age and sex, we added these variables as covariates in the regression models to test for main effects and interactions with the predictor variable.

## Results

### Multiparametric Biophysical Gradients in the Putamen

Our multiparametric analysis along the AP axis of the putamen has revealed distinct spatial variations in several biophysical parameters. Employing linear mixed-effects models (LMMs), we identified gradients of relaxation rates R1, R2, and R2*, WF, QSM, MTsat, and dMRI parameters MD and FA. These findings, depicted in Figure 1 and further detailed in Supplementary Table S1, align with previous research, notably replicating findings in WF, R1, and R2* and extend it to additional parameters^5^.

**Figure 1.**
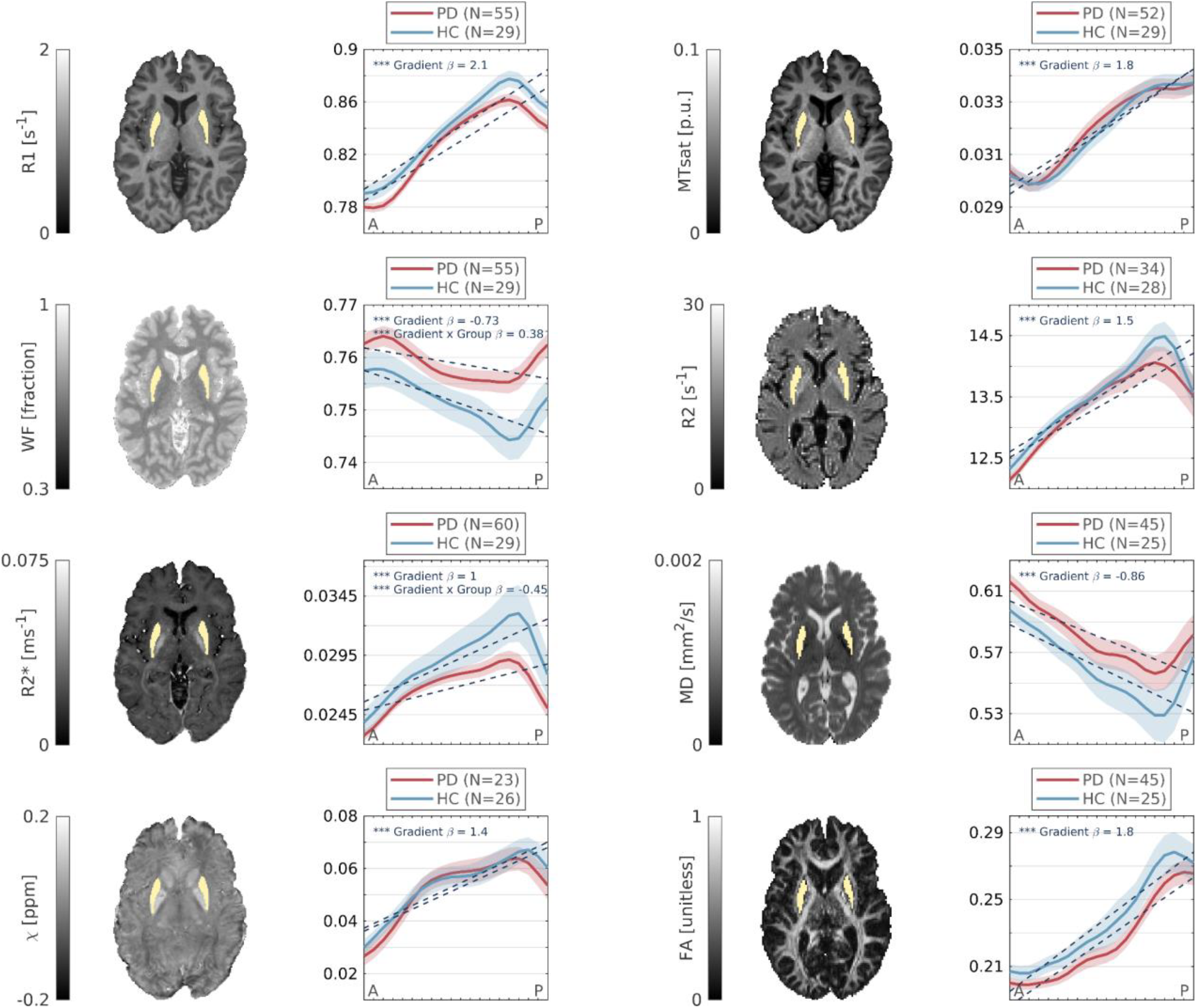
Putamen AP multiparametric gradients uncover local biophysical changes in PD. Spatial profiles of various biophysical mappings along the AP axis of the putamen uncover distinct and statistically significant gradients, as well as interactions with PD (Linear Mixed-effect models). Gradients of R1, WF, R2*, QSM (chi), MTsat, R2, MD and FA shown are averaged across the left and right putamen. Shaded area represents ± 1 SEM. Dashed lines represent the model’s linear fit. Standardized coefficient estimates (β) of spatial effect from anterior to posterior and PD group interaction are shown. * p < 0.01; ** p < 0.0001, *** p < 0.000001. Axial brain slices of a representative healthy subject are shown for each parameter, with bilateral putamen masks.

Notably, R1, R2, R2*, susceptibility, MTsat and FA showed generally common spatial patterns with values increasing towards posterior regions, and some decreases at the most posterior positions. WF and MD exhibited decreasing gradients toward posterior positions, followed by an increase at the posterior-most positions. This inverse behavior of WF and MD, compared to other parameters, is expected due to their opposing dependency on the tissue water content (see Discussion). Despite observed differences, all gradients were spatially intercorrelated, as depicted in Supplementary Figure S1.

To assess the nature of the observed spatial nonlinearities, we further expanded our analysis to include a nonlinear approach. Specifically, we repeated our LMM analysis while incorporating a quadratic Position term. Interestingly, this approach yielded statistically significant quadratic change in all MRI parameters, suggesting a non-linear aspect to their spatial variations (Supplementary Figure S2). While the quadratic term adds a layer of complexity that captures more nuanced spatial dynamics, we have opted to focus our interpretation and discussion on the linear approach, which is grounded in its simplicity and interpretability, and provides a balance between analytical rigor and practical understandability of later Group analyses.

### Microstructural Gradient Changes in Parkinson’s Disease

Intriguingly, the WF and R2* linear gradients significantly differed between PD patients and HC. This was evidenced by interaction effects between the Research Group and Spatial Position coefficients in the LMM analysis (Figure 1B, C, and Supplementary Table S1).

To further elucidate the gradient changes found in PD, we conducted post-hoc analyses that, in both WF and R2*, localized the maximal PD-related differences in the posterior putamen (PP). However, spatially dependent group differences, assessed via two-sample t-tests, were significant only in WF (Figure 2). Notably, traditional volumetric and median value analyses of the entire putamen failed to detect these differences, as shown in Supplementary Figure S3. This supports previous findings of PP alterations in PD^3–5^ and underscore the value of localized gradient analysis in revealing subtle microstructural changes.

**Figure 2.**
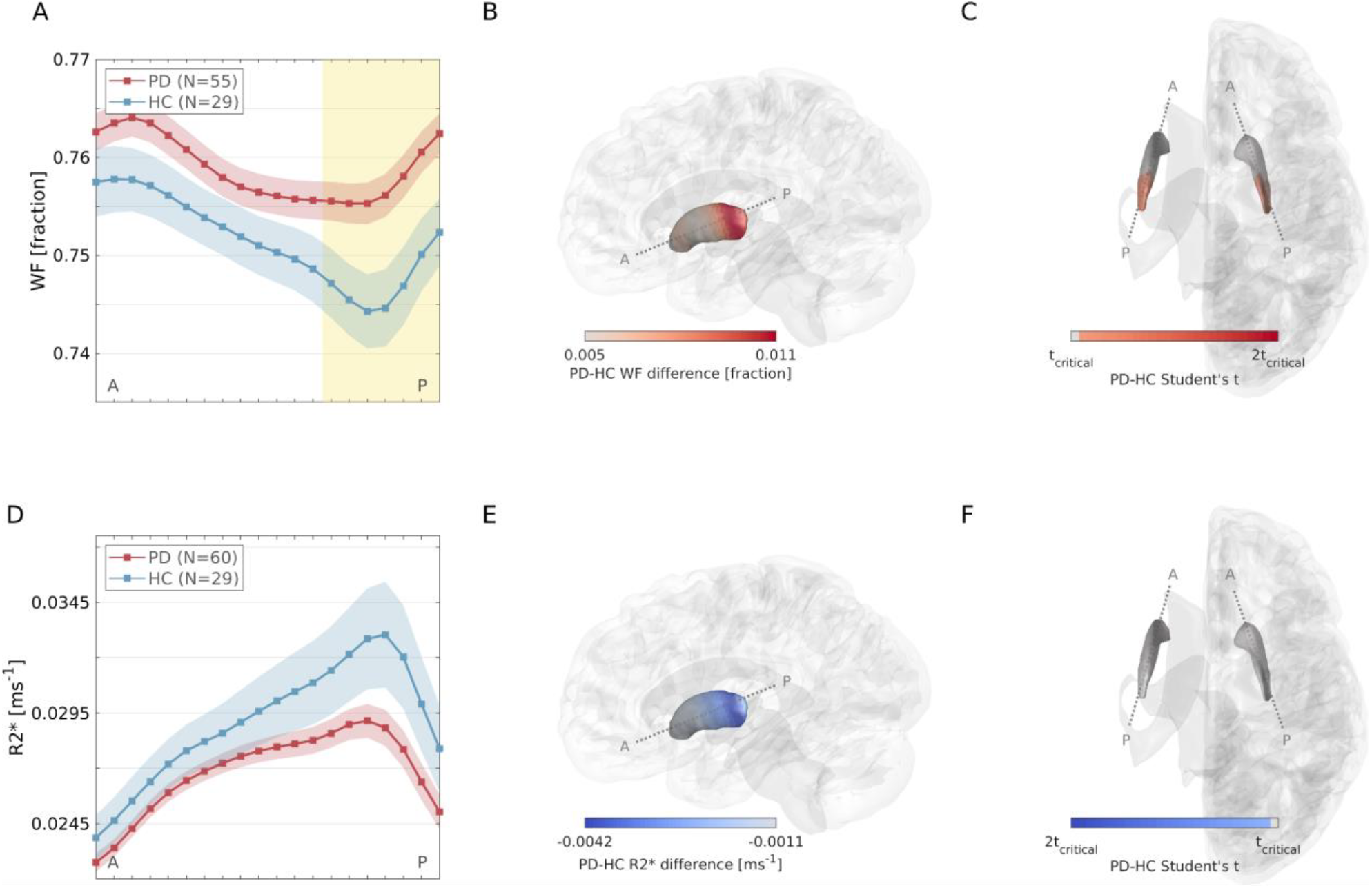
Local WF and R2* changes in PD. (A) Mean WF gradients in PD and HC. Shaded area represents ± 1 SEM. Yellow patch represents significant differences between groups, tested position-wise with two-sample t-tests (corrected p < .05). (B) Visualization of WF group differences in the putamen. PD-HC difference is positive (higher WF in PD, visualized in red). (C) Visualization of two-sample t-test significance. Insignificant values (below t critical) are visualized in gray. (D-F) Same analysis as in A-C, for R2*. PD-HC difference is negative (lower R2* in PD, visualized in blue), although statistically insignificant (absolute t values below absolute t critical).

Accounting for Age and Sex as covariates for the Research Group did not alter the significance of all observed effects. Nevertheless, a significant main effect of Sex was evident in R2* (Male *β* = -0.9 *p* = 0.007). These covariates exhibited no interaction with the Research Group, indicating that the PD-related effects found were independent of Age and Sex.

### Motor Symptom Laterality and Microstructural Changes

MRI-detected PP Alterations in PD patients have been previously linked to the body-side laterality of motor symptoms, assessed using MDS-UPDRS-III^5^. To shed light on the biophysical correlates of these PD characteristics, we tested the hypothesis that they are associated with changes in either water content (atrophy) or iron concentration. Specifically, given the PD-related gradient change we found in WF and R2*, we examined the relationships of these parameters in the PP with the patients’ motor body-side laterality.

First, in patients with motor sign asymmetries (N = 45; See Methods), we compared WF and R2* gradients between the most-affected side (MAS) and less-affected side (LAS). Our findings showed a significantly higher WF in the MAS, suggesting increased tissue-water content in the affected side. This effect was spatially dependent, predominantly located in the PP, albeit slightly more anterior than regions differing from healthy controls (Figure 3A). In contrast, no significant differences in R2* gradients were observed between MAS and LAS (Supplementary Figure S4A).

**Figure 3.**
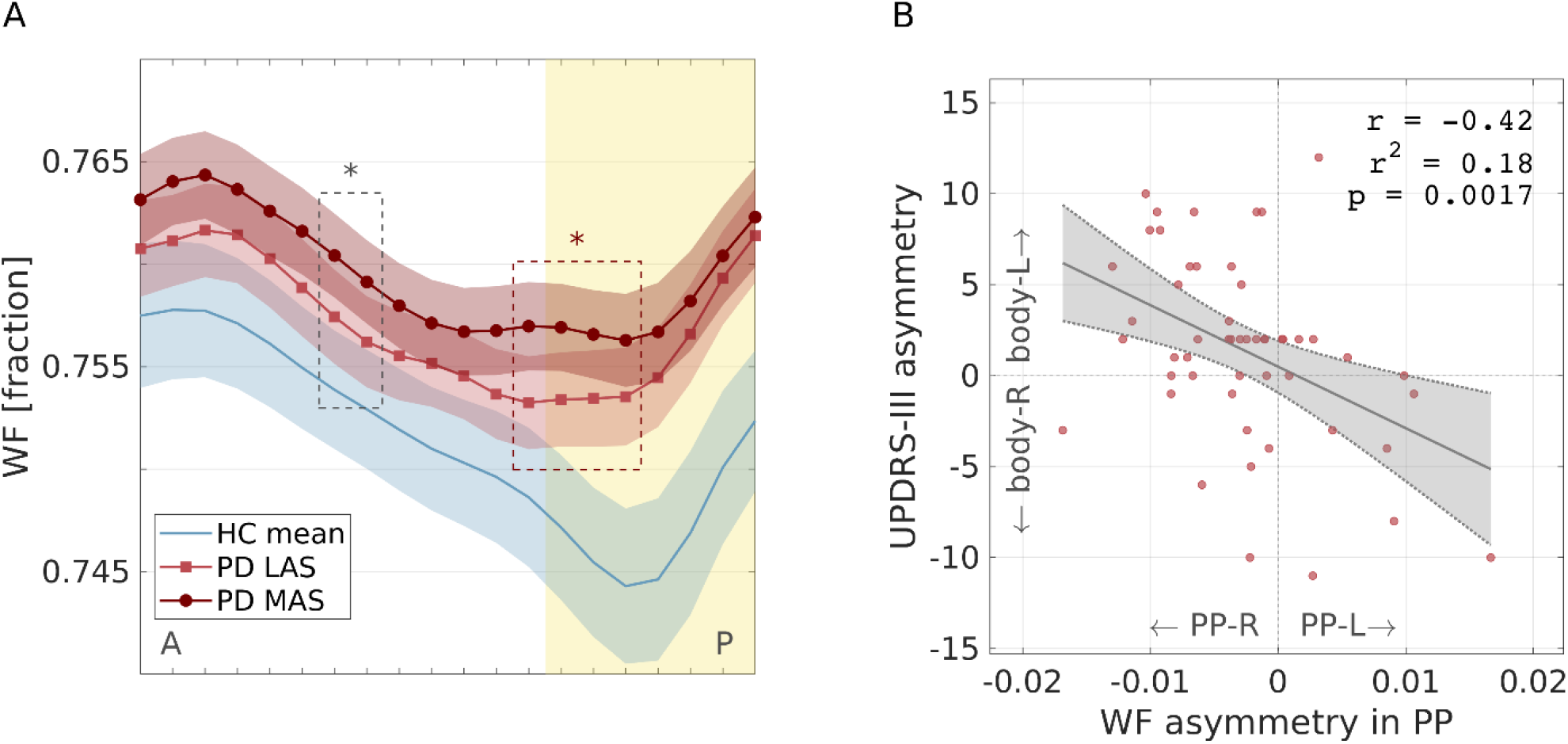
WF asymmetry in PP associated with contralateral motor signs. **(A)** WF gradients in the putamen of patients with motor body-side predominance (N=46 having motor asymmetry) and healthy controls. PD gradients are grouped into LAS and MAS, based on patients’ contralateral motor laterality. Yellow patch represents difference from HC (see Fig. 2). Asterisks represent significant differences between LAS and MAS. Posterior significant positions (marked in red rectangle) are used for computing the PP WF asymmetry displayed in panel B. **(B)** WF asymmetry between left and right PP in PD patients (N=52 with available UPDRS III data) is significantly correlated with contralateral motor signs asymmetry. PP positions used in this analysis are those marked in red rectangle in panel A. Pearson’s r = -0.42; r^2^ = 0.18; p = 0.0018

Additionally, we explored the relationship between inter-hemispheric asymmetry in the PP and patients’ motor asymmetry score. We found a significant correlation between WF asymmetry and motor symptom asymmetry, with higher WF associated with contralateral symptoms (Figure 3B). In contrast, R2* asymmetry did not show a significant correlation with motor asymmetry (Supplementary Figure S4B). These findings may suggest that motor symptom laterality in PD is primarily associated with increased water fraction in the contralateral PP rather than variations in iron concentration.

However, the absence of observable differences in R2*, aside from gradient change, might also be attributed to the inherently higher noise levels associated with R2* measurements. The average inter-subject coefficient of variation (CV) in the R2* was 23.3%, compared to 2.2% in WF. This high inter-subject variance in R2* is potentially obscuring subtle changes that are observed in WF.

Finally, we sought to investigate PD-related changes in tissue macromolecular composition and in iron environment homeostasis, rather than changes in quantities of tissue and iron. Filo et al.^38,39^ introduced the Tissue and Iron Relaxivity frameworks, showing that the linear dependencies of R1 on the non-water fraction (dR1/d(1-WF)) and on R2* (dR1/dR2*), are sensitive to the macromolecular composition and iron homeostasis, respectively. We calculated these dependencies in the PP and tested for differences between PD and HC. Our analysis did not reveal statistically significant differences between the groups (two-sample t-tests, *p* > 0.16), thereby failing to indicate changes in macromolecular compositions or in iron homeostasis. Consequently, this supports the conclusion that the effect found in WF is more likely due to a reduction in macromolecular content, indicative of tissue atrophy, rather than alterations in lipid or iron composition.

## Discussion

Our study leveraged qMRI and spatial gradient approach to investigate the putamen, revealing deep brain biophysical changes in both healthy individuals and PD patients. These investigations have provided several critical insights. Firstly, our findings confirmed the existence of microstructural gradients within the putamen, corroborating the concept of striatal gradients and demonstrating the capability of MRI to capture these spatial variations in a living brain. This supports the notion that the putamen’s structure varies systematically along its AP axis, which has implications for understanding its functional organization and connectivity in both health and disease.

Secondly, we identified localized neurodegenerative changes in the PP, predominantly related to water increase in PD patients. The absence of volumetric differences in our data supports the conclusion that this finding reflects micro- or mesoscale, rather than macroscale, alterations in the PP. This change in water fraction, at the expense of the lipid and macromolecular tissue content, can be attributed to lower non-water cellular tissue^38^, as well as to neuroinflammation processes which involve elevation of extracellular water content^40^. This finding is particularly significant as it aligns with the hypothesis that PD pathology initiates in specific brain regions and progresses in a spatially structured manner. Our results add to the growing body of evidence suggesting that the PP may be a critical site for early PD-related changes, further emphasizing the importance of spatially resolved analyses in detecting and understanding the progression of neurodegenerative diseases.

Lastly, our study established a link between the asymmetry in water increase within the PP and the lateralization of motor symptoms in PD patients, corroborating earlier results in clinical MRI^5^, PET and SPECT^4^. This correlation underscores the functional relevance of the observed microstructural alterations, indicating that the side of the brain exhibiting more severe tissue loss is associated with more pronounced motor impairment on the opposite side of the body. Such findings highlight the potential of qMRI to provide biomarkers that not only reflect the underlying neuropathology of PD but also correlate with clinical manifestations of the disease, offering a more integrated view of brain structure and function.

While all qMRI metrics showed spatial correlation, only select parameters, namely WF and R2*, demonstrated PD-related alterations. These changes suggest potential markers for tissue atrophy (indicated by increased tissue-water) and possibly reduced iron content, as implied by R2* decrease. However, the R2* findings warrant cautious interpretation. We observed a spatially dependent decrease in R2* in PD, characterized by a less pronounced gradient slope. Additionally, although not reaching statistical significance in two-sample t-tests, mean R2* values in the PP and in the entire putamen showed lower trends in PD patients. Contrary to expectations of increased R2* due to iron deposition in PD, the literature presents mixed results, with several studies not finding significant differences in R2* values between PD and HC in the PP^41,42^. Importantly, while R2* is commonly correlated with iron content, it is also influenced by other factors such as myelin content, fiber orientation, calcium, tissue integrity, vascularization, and water content^43^. This poses a challenge in interpreting the biological sources of R2* changes and their direct correlation with PD pathology. The relatively high coefficient of variation (CV) in R2* suggests greater noise, potentially masking significant changes and leading to less definitive results.

In our study, the linear modeling effectively uncovered microstructural gradients along the anterior-posterior axis of the putamen. In addition, these linear gradients highlighted PD-related changes, consistent with previous literature on PD neurodegeneration. Specifically, linear gradients of WF and R2* provided clear evidence of group differences in the PP, possibly reflecting the gradual, posteriorly-initiating loss of dopaminergic terminals and alterations in tissue content. The strength of linear modeling is its relative simplicity and interpretability. Nonetheless, our supplementary quadratic analysis unveiled additional insights, revealing a more complex spatial pattern of changes, not captured by the linear model.

The quadratic analysis revealed nonlinear spatial changes along the putamen’s anterior-posterior axis, indicating that non-linear variations in microstructural properties are inherent to the putamen’s anatomy. The significant interaction between the research group and the quadratic spatial term suggests differences in these inherent non-linearities between PD and HC. Specifically, interactions between the quadratic term of the AP position and PD were found in R1, WF, susceptibility and MTsat.

Thus, while the linear approach provides a straightforward depiction of overall trends, the quadratic analysis enriches our understanding by identifying localized, non-linear changes that could be pivotal in understanding the nuanced mechanisms of PD progression. This emphasizes the importance of considering both linear and non-linear dynamics when investigating the microstructural changes in PD, as it could uncover specific pathological mechanisms that are not apparent in a purely linear analysis.

### Limitations

Our study faces several limitations that merit consideration. Firstly, the comprehensive qMRI battery required for detailed microstructural analysis necessitated long scanning sessions, approximately 1.5 hours, posing challenges for subjects. This extensive scan time led to variability in the completion rates of different qMRI parameters, as well as motion artifacts, resulting in varied sample sizes across metrics and potentially impacting the robustness of our findings. Future studies could benefit from employing shorter imaging protocols, such as Magnetic Resonance Fingerprinting (MRF)^44,45^, to reduce scan time and minimize the potential for motion artifacts, thereby improving patient comfort and data consistency.

The heterogeneous nature of our cohort, encompassing a wide range of disease stages, ages, and cognitive statuses, introduces additional variability into our analysis. This diversity, while offering a broad perspective on PD, may introduce confounding factors that may interact with our results.

Furthermore, the resolution discrepancies among qMRI metrics pose challenges for direct comparisons. While most qMRI maps were acquired with a resolution of 1 mm, the DTI and R2 measurements were obtained at coarser resolutions of 1.5 mm and 2 mm, respectively. This variance in resolution limits the statistical comparability between parameters and may affect the interpretation of spatial gradients and their pathological significance.

In conclusion, our study leverages multiparametric qMRI to elucidate microstructural changes in PD, offering new insights into its neuropathology and clinical features. By providing a detailed analysis of the putamen’s biophysical properties, we have identified potential biomarkers for disease detection and manifestation. Our findings underscore the importance of a comprehensive imaging approach to capture subtle, spatially dependent biological alterations in PD. The ability to quantitatively measure microstructural changes in a progressive disease like PD, presents opportunities for monitoring disease progression during studies evaluating potential disease-modifying therapies. Future research may benefit from exploring the heterogeneity within PD populations, ultimately contributing to more personalized and effective diagnostic and therapeutic strategies.

## Acknowledgments

We thank S. Filo and S. Moskovich for their critical reading of the manuscript and their useful comments.

