## Supplementary material for "Multiparametric quantitative MRI uncovers putamen microstructural changes in Parkinson’s Disease": Supplemetary Materials

By Drori et. al (2024)

### Supplementary Table S1

***Table S1 – Linear Mixed Effects Models (LMM) Results**. Results of the LMM with linear Position term and no covariates. Coefficients are in the native response units (MRI parameter units). Β coefficients are standardized for comparison between model responses. Position effects represent the global change from anterior to posterior (regardless of number of nodes along the axis)*

| **R1** | Coefficient (95% CI) | *β* (95% CI) | t | DF | p | p_FDR |
|---|---|---|---|---|---|---|
| ClinicalGroup_PD | -0.009 (-0.022, 0.005) | -0.201 (-0.506, 0.105) | -1.29 | 1680 | 0.197 | 0.197 |
| Position | 0.091 (0.086, 0.096) | 2.090 (1.970, 2.200) | 35.5 | 1680 | 3.00E-206 | 1.00E-205 |
| ClinicalGroup_PD:Position | -0.005 (-0.011, 0.001) | -0.110 (-0.253, 0.032) | -1.52 | 1680 | 0.13 | 0.173 |
| **WF** | Coefficient (95% CI) | *β* (95% CI) | t | DF | p | p_FDR |
| ClinicalGroup_PD | 0.004 (-0.002, 0.011) | 0.257 (-0.149, 0.663) | 1.24 | 1680 | 0.214 | 0.286 |
| Position | -0.012 (-0.014, -0.010) | -0.726 (-0.828, -0.624) | -13.9 | 1680 | 1.00E-41 | 4.00E-41 |
| ClinicalGroup_PD:Position | 0.006 (0.004, 0.008) | 0.375 (0.249, 0.502) | 5.82 | 1680 | 7.00E-09 | 1.00E-08 |
| **R2*** | Coefficient (95% CI) | *β* (95% CI) | t | DF | p | p_FDR |
| ClinicalGroup_PD | -0.001 (-0.003, 0.002) | -0.104 (-0.486, 0.278) | -0.533 | 1780 | 0.594 | 0.594 |
| Position | 0.007 (0.006, 0.008) | 1.030 (0.903, 1.160) | 15.9 | 1780 | 2.00E-53 | 6.00E-53 |
| ClinicalGroup_PD:Position | -0.003 (-0.004, -0.002) | -0.455 (-0.609, -0.301) | -5.78 | 1780 | 9.00E-09 | 2.00E-08 |
| **susceptibility** | Coefficient (95% CI) | *β* (95% CI) | t | DF | p | p_FDR |
| ClinicalGroup_PD | -0.001 (-0.011, 0.009) | -0.050 (-0.482, 0.381) | -0.229 | 976 | 0.819 | 0.819 |
| Position | 0.033 (0.029, 0.036) | 1.410 (1.250, 1.560) | 18.3 | 976 | 2.00E-64 | 7.00E-64 |
| ClinicalGroup_PD:Position | -0.001 (-0.006, 0.004) | -0.037 (-0.257, 0.183) | -0.327 | 976 | 0.743 | 0.819 |
| **Mtsat** | Coefficient (95% CI) | *β* (95% CI) | t | DF | p | p_FDR |
| ClinicalGroup_PD | 0.000 (-0.001, 0.001) | 0.109 (-0.248, 0.466) | 0.598 | 1620 | 0.55 | 0.55 |
| Position | 0.005 (0.004, 0.005) | 1.770 (1.660, 1.870) | 34.1 | 1620 | 3.00E-192 | 1.00E-191 |

| | | | | | | |
|---|---|---|---|---|---|---|
| ClinicalGroup_PD:Position | -0.000 (-0.001, 0.000) | -0.109 (-0.236, 0.018) | -1.68 | 1620 | 0.092 | 0.123 |
| **R2** | Coefficient (95% CI) | *β* (95% CI) | t | DF | p | p_FDR |
| ClinicalGroup_PD | -0.091 (-0.536, 0.355) | -0.075 (-0.443, 0.293) | -0.399 | 1240 | 0.69 | 0.69 |
| Position | 1.860 (1.670, 2.040) | 1.530 (1.380, 1.690) | 19.6 | 1240 | 5.00E-75 | 2.00E-74 |
| ClinicalGroup_PD:Position | -0.157 (-0.407, 0.093) | -0.130 (-0.336, 0.077) | -1.23 | 1240 | 0.219 | 0.291 |
| **MD** | Coefficient (95% CI) | *β* (95% CI) | t | DF | p | p_FDR |
| ClinicalGroup_PD | 0.000 (-0.000, 0.000) | 0.231 (-0.175, 0.637) | 1.11 | 1400 | 0.265 | 0.265 |
| Position | -0.000 (-0.000, -0.000) | -0.858 (-1.010, -0.702) | -10.8 | 1400 | 3.00E-26 | 1.00E-25 |
| ClinicalGroup_PD:Position | 0.000 (-0.000, 0.000) | 0.138 (-0.055, 0.332) | 1.4 | 1400 | 0.162 | 0.215 |
| **FA** | Coefficient (95% CI) | *β* (95% CI) | t | DF | p | p_FDR |
| ClinicalGroup_PD | -0.009 (-0.024, 0.007) | -0.187 (-0.521, 0.146) | -1.1 | 1400 | 0.27 | 0.27 |
| Position | 0.083 (0.076, 0.090) | 1.830 (1.680, 1.990) | 22.8 | 1400 | 2.00E-98 | 1.00E-97 |
| ClinicalGroup_PD:Position | -0.007 (-0.015, 0.002) | -0.145 (-0.342, 0.051) | -1.45 | 1400 | 0.147 | 0.195 |

### Supplementary Figure S1

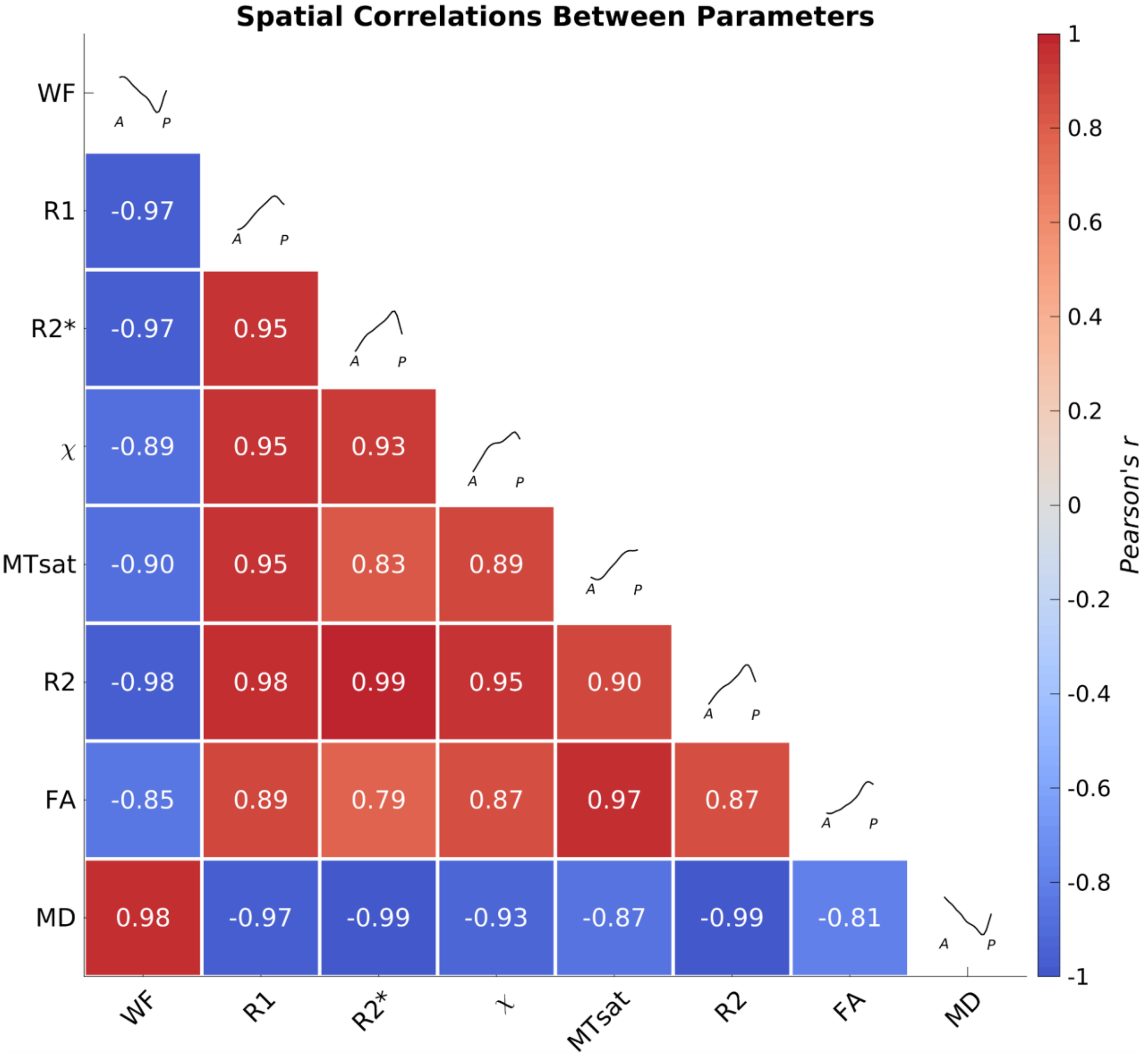

***Figure S1 – Multiparameter gradients along the putamen's AP axis are spatially correlated.*** *Correlation matrix summarizes putamen AP gradient spatial covariance between each two MRI parameters. All putamen gradients are spatially intercorrelated. While R1, R2*, susceptibility, Mtsat, R2, and FA are positively correlated, they are negatively correlated with WF and MD. The mean putamen gradients of healthy controls are presented on the diagonal for each parameter. Qualitatively similar results were found in PD patients.*

### Supplementary Figure S2

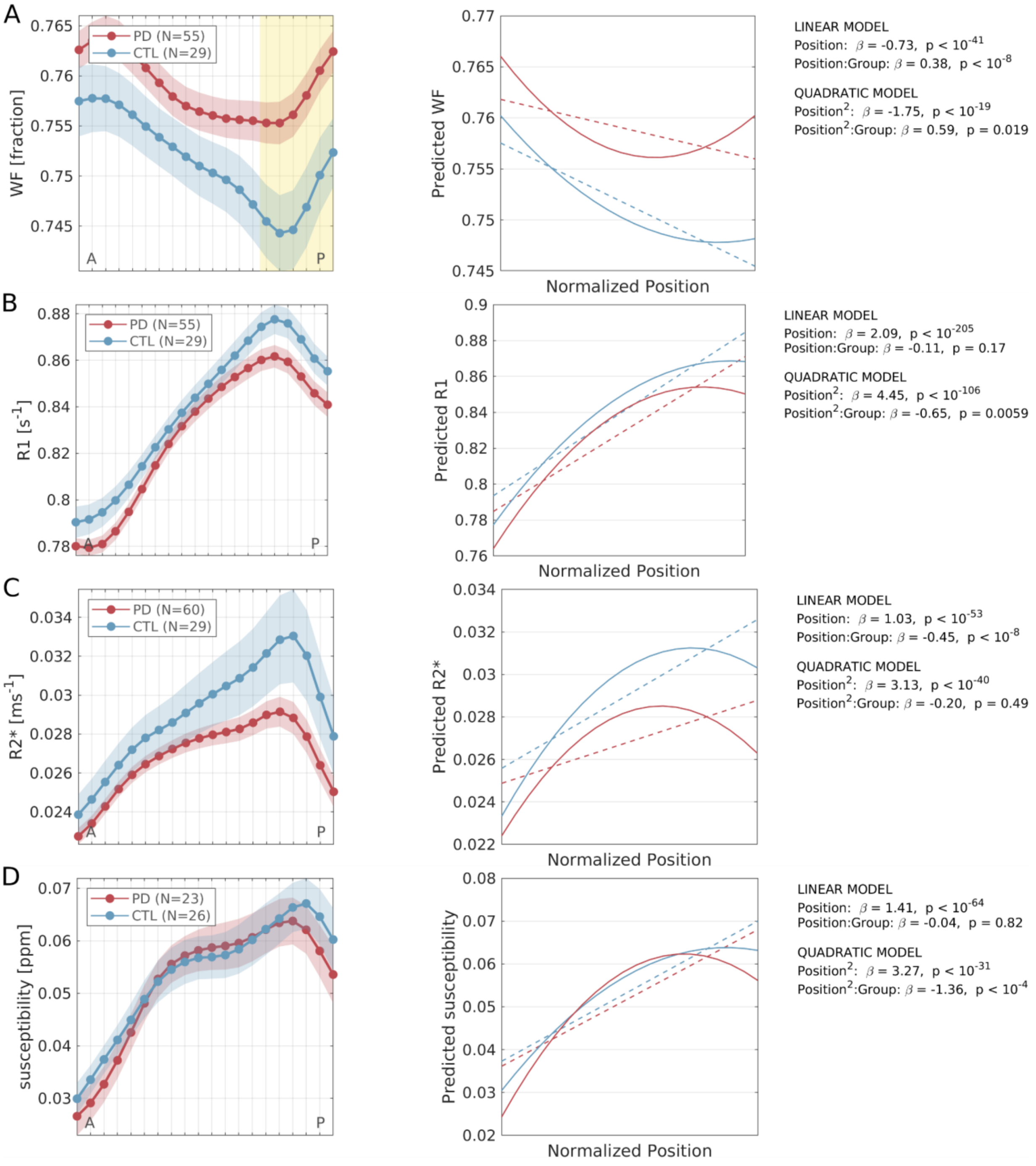

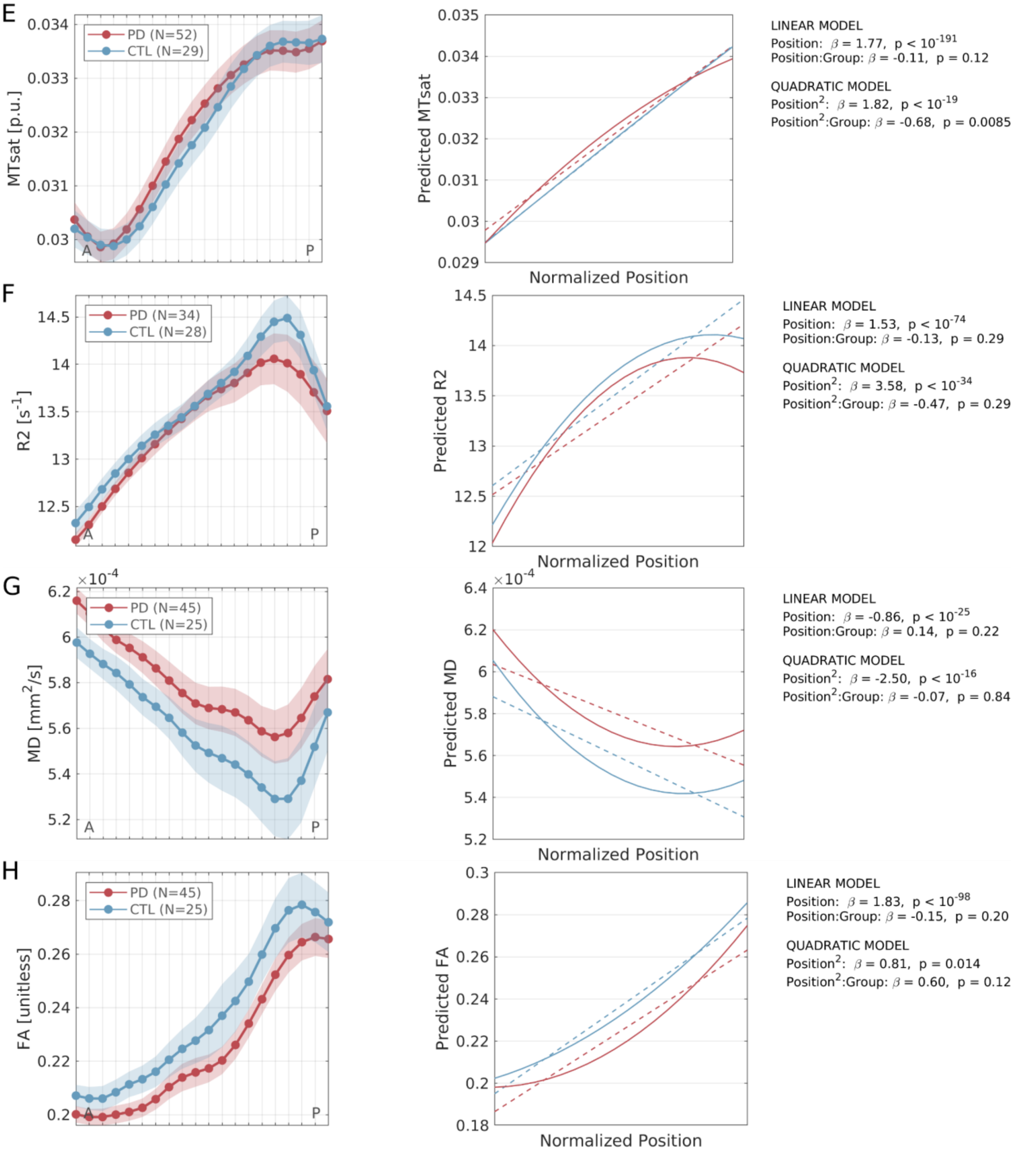

***Figure S2 – Quadratic analysis of multiparametric putamen AP gradients.*** *Left panels – putamen AP profiles of PD and HC (as in Fig. 1). Middle panels – linear (dashed lines) and quadratic (solid lines) model fits. Right panels – the standardized beta coefficients and p-values of the linear and quadratic spatial Position terms and their interactions with the Group term. The quadratic terms of all models are significant, suggesting nonlinear spatial change. The interaction of quadratic term with the research group is significant in the WF, R1, Mtsat and susceptibility models, suggesting subtle changes between groups in the quadratic fit.*

### Supplementary Figure S3

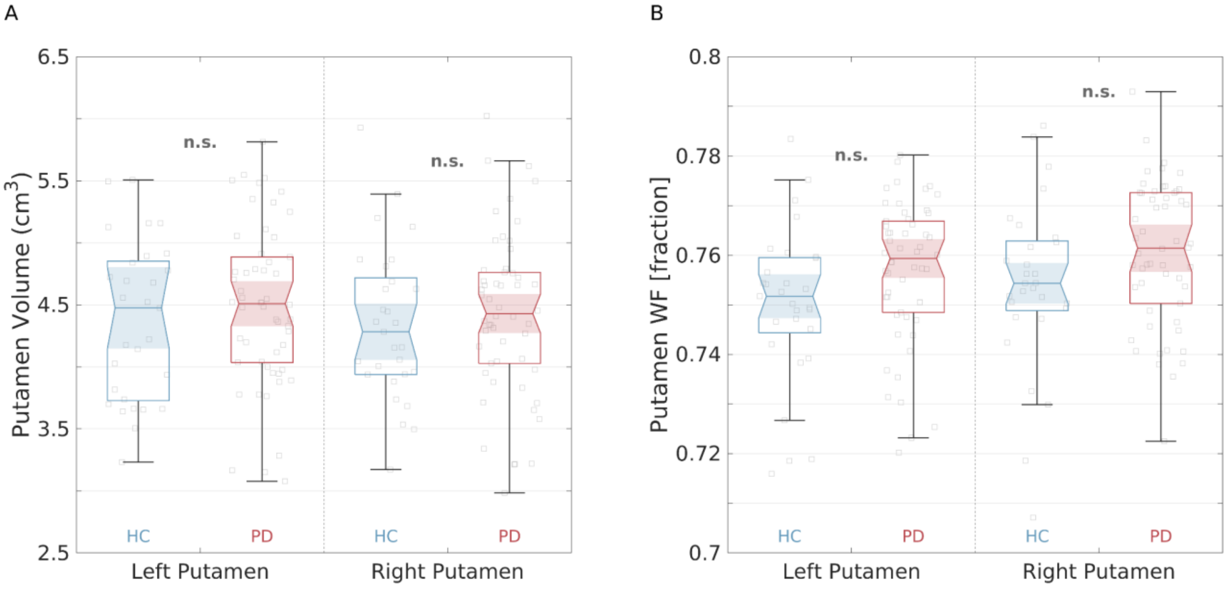

***Figure S3 - No group differences in whole-ROI analysis.*** *PD (N=54) and HC (N=29) did not significantly differ in (A) whole-putamen volume or (B) whole-putamen WF. Similar results (n.s.) was obtained for all MRI parameters (R1, R2*, susceptibility, MTsat, R2, MD and FA).*

### Supplementary Figure S4

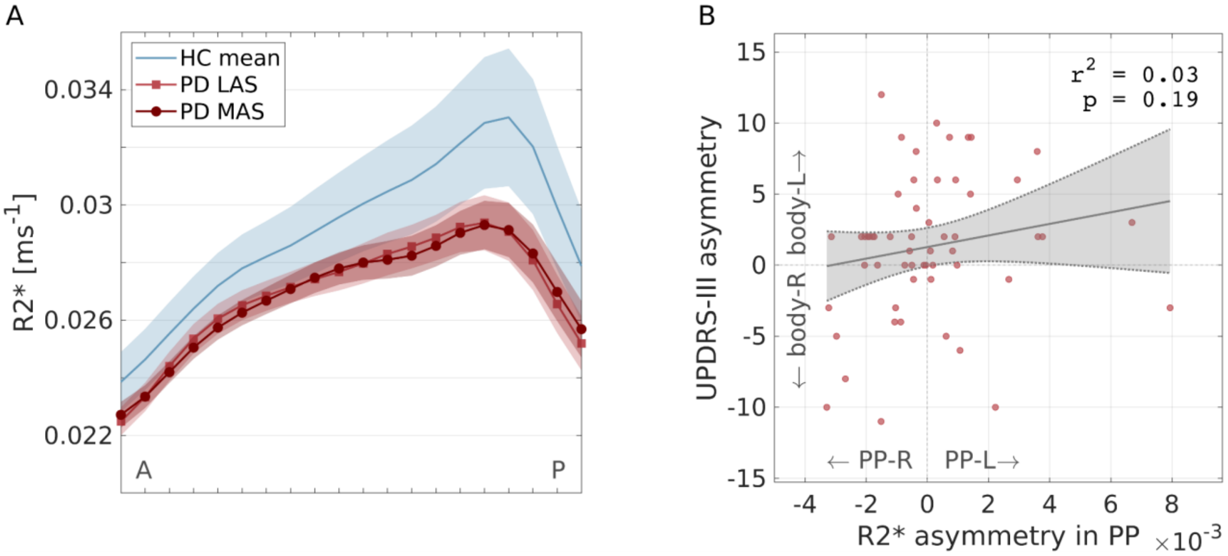

***Figure S4 – R2* in PP is not correlated with motor asymmetry. (A)*** *R2* gradients in the putamen of patients with motor side predominance (N=45 having motor asymmetry) and healthy controls. PD gradients are grouped into LAS and MAS, based on patients' contralateral motor laterality. No significant differences between LAS and MAS were found.* ***(B)*** *R2* asymmetry between left and right PP in PD patients (N=51 with available UPDRS III data) is not correlated with contralateral motor signs asymmetry. PP positions used in this analysis are those used in Figure 3. Pearson's r = -0.42; r2 = 0.18; p = 0.0018*
